# sc-SPLASH provides ultra-efficient reference-free discovery in barcoded single-cell sequencing

**DOI:** 10.1101/2024.12.24.630263

**Authors:** Roozbeh Dehghannasiri, Marek Kokot, Alexander L. Starr, Jamie Maziarz, Tal Gordon, Serena Y. Tan, Peter L. Wang, Ayelet Voskoboynik, Jacob M. Musser, Sebastian Deorowicz, Julia Salzman

## Abstract

Typical high-throughput single-cell RNA-sequencing (scRNA-seq) analyses are primarily conducted by (pseudo)alignment, through the lens of annotated gene models, and aimed at detecting differential gene expression. This misses diversity generated by other mechanisms that diversify the transcriptome such as splicing and V(D)J recombination, and is blind to sequences missing from imperfect reference genomes. Here, we present sc-SPLASH, a highly efficient pipeline that extends our SPLASH framework for statistics-first, reference-free discovery to barcoded scRNA-seq (10x Chromium) and spatial transcriptomics (10x Visium); we also provide its optimized module for preprocessing and *k*-mer counting in barcoded data, BKC, as a standalone tool. sc-SPLASH rediscovers known biology including V(D)J recombination and cell-type-specific alternative splicing in human and trans-splicing in tunicate (*Ciona*) and when applied to spatial datasets, detects sequence variation including tumor-specific somatic mutation. In sponge (*Spongilla*) and tunicate (*Ciona*), we uncover secreted repeat proteins expressed in immune-type cells and regulated during development; the sponge genes were absent from the reference assembly. sc-SPLASH provides a powerful alternative tool for exploring transcriptomes that is applicable to the breadth of life’s diversity.

## Main text

Single-cell RNA-sequencing (scRNA-seq) has great potential to reveal cellular identity and phenotypes. In particular, barcoded droplet-based scRNA-seq (such as 10x Genomics Chromium) can process thousands of cells per run. Although the resulting data contains information on many molecular mechanisms such as RNA alternative splicing, V(D)J rearrangement, and RNA editing among others, most scRNA-seq studies remain focused on gene expression due to computational and statistical challenges and lack of alternative tools. While a few computational tools have emerged to facilitate single-cell analysis beyond gene expression (Olivieri et al. 2021; Olivieri, Dehghannasiri, and Salzman 2022; Xiang et al. 2024; Cuddleston et al. 2022; Sturm et al. 2020; Borcherding, Bormann, and Kraus 2020; Meyer et al. 2022; Gao et al. 2021; Patrick et al. 2020), their adoption in scRNA-seq studies remains limited. These tools are fragmented, fine-tuned to detect only a specific event, and their installation and execution typically require extensive bioinformatic expertise. On the other hand, these tools are prone to biases and blind spots due to their reliance on the computationally intensive alignment of noisy sequencing reads to reference genome, which could lead to the overlooking of key biological mechanisms not represented in the reference genome (Chaung et al. 2023), a challenge particularly common in non-model organisms which often lack high-quality reference genomes (Ungaro et al. 2017).

To address these challenges, we leverage the recent SPLASH framework (Chaung et al. 2023; Kokot et al. 2024), which performs statistical inference directly on raw sequencing reads. SPLASH bypasses sequence alignment and employs a *k*-mer-based approach to detect sample-specific sequence diversity variation caused by myriad mechanisms such as alternative splicing, RNA editing, V(DJ) recombination, etc. SPLASH first parses reads from all input samples to identify specific *k*-mers (called *anchors*) that are followed by a set of diverse downstream *k*-mers (called *targets*) and then utilizes a statistical hypothesis test to assign a closed-form p-value where anchors with significant p-values exhibit sample-dependent target distribution. SPLASH has uncovered novel biological insights, both from bulk RNA-seq and plate-based (Smart-seq2) scRNA-seq (Chaung et al. 2023; Kokot et al. 2024; Dehghannasiri et al. 2022). While plate-based scRNA-seq typically accesses hundreds of cells, barcoded droplet scRNA-seq can process orders of magnitude more; thus extension of SPLASH to barcoded scRNA-seq requires dealing with its more complex data structure and performance challenges.

Here, we introduce sc-SPLASH, an ultra-efficient easy-to-use pipeline for analyzing transcriptomic complexity in barcoded scRNA-seq, as an alternative to reference-based, gene expression-centric approaches. It performs statistical inference directly on raw sequencing reads to detect regulated sequence diversity and performs versatile downstream analyses all with just a single-line command. sc-SPLASH in built on SPLASH2 (Kokot et al. 2024), our work for bulk RNA-Seq analysis, and similar to SPLASH2 has three core stages (Figure 1A): 1) parsing reads and extracting anchor-targets; 2) merging anchor-target pairs and their counts across cells, and computing p-values; and 3) performing multiple testing correction. Barcoded scRNA-seq data is more complex because thousands of cells are pooled into a single library. Each read is associated with a cell barcode and a unique molecular identifier (UMI), observed with sequencing error. Thus, we completely redesigned Stage 1 relative to SPLASH2 and particularly developed BKC, an optimized tool for *k*-mer counting in barcoded data(analogous to KMC for bulk sequencing data (Kokot, Dlugosz, and Deorowicz 2017)). Like other 10x preprocessing tools, BKC extracts cell barcodes and performs UMI deduplication (Methods), but it is considerably faster, running 43-fold faster than the widely used UMI-tools package (Smith, Heger, and Sudbery 2017)(Methods). BKC then separately parses reads assigned to each extracted cell to obtain either *k*-mer or paired *k*-mer (anchor-target) counts. Unlike other preprocessing tools, BKC also includes a rapid step to filter sequencing artifacts due to Illumina adapters or other user-provided contaminants through the implementation of hash-tables and Bloom filters. We provide BKC as a standalone tool, as we anticipate it could be valuable for developing other pipelines.

**Figure 1.**
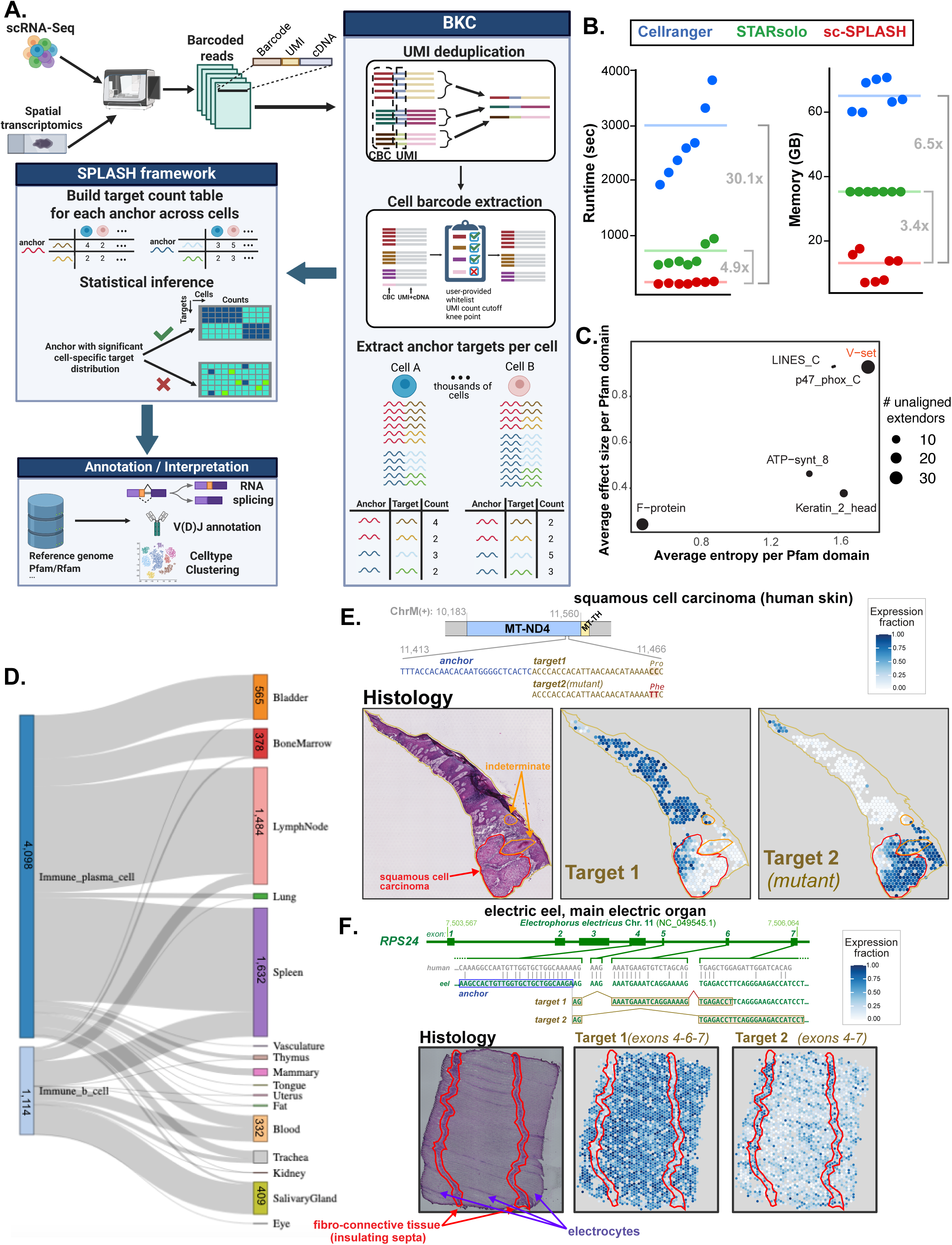
sc-SPLASH pipeline overview and analysis of human V(D)J rearrangement and spatial transcriptomics (Visium). **A.** Overview of sc-SPLASH pipeline including preprocessing of 10x scRNA-seq data (cell barcode extraction and UMI deduplication) and performing anchor/target counting through the BKC module and then performing statistical analysis to identify anchors (constant sequences) followed by a diverse set of target sequences with single-cell dependent distribution. **B.** Comparison of sc-SPLASH with Cellranger and STARsolo, two state-of-the-art 10x processing tools indicates much higher efficiency for scSPLASH. Test samples are from Tabula Sapiens dataset and are ordered by size. **C.** Pfam analysis on unaligned extendors suggests that the immunoglobulin variable domain (V-set) has the highest number of unaligned extendors and also highest average entropy compared to other Pfam domains. **D.** Distribution of cell types and tissues containing in-frame V(D)J transcripts identified by sc-SPLASH+IgBLAST. **E.** Detection of a tumor-associated a double somatic mutation in gene *MT-ND4* in squamous cell carcinoma Visium data by sc-SPLASH where Target 2 corresponding to the mutation has higher fraction in carcinoma cells (marked by red boundary). **F.** Spatially-regulated alternative splicing of *RPS24* detected in electric eel Visium data where electrocytes (purple arrows) include exon 6 and stromal cells in the insulating septa (red arrows) exclude exon 6. We also show the correspondence between the RPS24 nucleotide sequence in human and electric eel.

Stage 2 of sc-SPLASH was designed to address the increased memory requirements of 10x data as each record is larger, including a cell barcode in addition to the anchor, target, and sample ID, with many many such records loaded into memory simultaneously. For each anchor, a contingency table, containing target counts across cells is constructed. With thousands of cells in a scRNA-seq experiment, these contingency tables can become quite large. We leveraged a sparse matrix representation and further engineering optimization to address this large memory demand. A p-value is computed for each contingency table (Baharav, Tse, and Salzman 2024) and multiple testing correction is performed in Stage 3, similar to SPLASH2. For anchors with significant p-values a number of post-processing analyses can be done to aid biological interpretation. If metadata exists, it can be used to identify metadata-dependent anchors (i.e., cell-type-specific anchors) using a supervised test based on an L1-regularized generalized linear model as implemented in GLMnet (Friedman, Hastie, and Tibshirani 2010) (Methods). sc-SPLASH can construct “extendors” by concatenating anchor-target pairs which can be aligned to a reference genome *post facto* to identify anchors due to splicing and single base-pair changes or even without a genome be queried for BLAST matches and Pfam protein domains.

We compared sc-SPLASH performance against two widely-used tools for 10x analysis: Cell Ranger (Zheng et al. 2017) and STARsolo (Kaminow, Yunusov, and Dobin 2021) (Methods). sc-SPLASH (up to calling significant anchors) was almost 5 times and 30 times faster than STARsolo and Cell Ranger, respectively, while also requiring considerably less memory than both (Figure 1B). This is particularly noteworthy as Cell Ranger and STARsolo handle only the initial sequence alignment and require other tools such as Seurat (Satija et al. 2015) or Scanpy (Wolf, Angerer, and Theis 2018) for downstream differential gene expression analysis. This demonstrates the high overhead of alignment-based approaches.

To benchmark sc-SPLASH, we applied it to over 400,000 human single cells from the Tabula Sapiens (TS) 10x dataset (Tabula Sapiens Consortium* et al. 2022). Using the supervised test with cell type metadata, we identified 555 genes with sequence diversity differing between cell types. Among these, *RPS24* and *MYL6* were found in most donor tissues (37 and 28, respectively), consistent with their known cell type-specific alternative splicing (Olivieri et al. 2021).

Pfam analysis of extendors for all anchors called by sc-SPLASH across TS dataset showed that those mapping to the variable domain of the immunoglobulin superfamily (V-set) had the highest entropy (2.16) and effect size (0.90) compared to those mapping to other Pfam domains, which is consistent with our previous work on plate-based scRNA-seq where anchors for antibody (Ig) and T-cell receptor (TCR) had a characteristically high diversity of targets (Chaung et al. 2023). Among unaligned extendors, V-set domain had the highest number of extendors (Figure 1C), suggesting the challenges of aligning V(D)J transcripts. We also identified a viral fusion protein domain (F-protein) in the heart cells from donor 12 and confirmed that sequencing data included human immunodeficiency virus transcripts (Supplement), highlighting the utility of not solely relying on genome alignment. (Figure 1C). We integrated sc-SPLASH with our recently developed method for assembling longer sequences (Henderson et al. 2024) and with IgBLAST (Ye et al., 2013) for V(D)J annotation (Methods). Across the TS 10x dataset, we detected 60,697 assembled V(D)J sequences, 46% of which were productive in-frame transcripts (Figure 1D). SPLASH may be a useful adjunct for identifying diversifying genes like Ig/TCR, especially in organisms with less well-defined references.

Spatially-resolved transcriptomics is an emerging technology for investigating RNA regulation in its natural tissue context. Particularly interesting are spatial methods such as 10x Visium, which are based on sequencing rather than hybridization and thus not restricted to prespecified sets of genes.

Although sc-SPLASH was developed for barcoded scRNA-seq, given that Visium has the same barcoded data format, sc-SPLASH can be readily applied to it. As a demonstration, sc-SPLASH was used on several Visium samples: human cutaneous squamous cell carcinoma (Ji et al. 2020), human fetal intestine (Fawkner-Corbett et al. 2021), and eel electric organ (Methods). In the squamous carcinoma tissue sample, the anchor with the highest effect size (0.757) revealed a double mutation CC to TT in the mitochondrial gene *MT-ND4* and its wild-type counterpart. The mutated version was predominantly (but not exclusively) expressed in the area of carcinoma (Figure 1E). This pattern is consistent with the mutation arising spontaneously in the lineage of the carcinoma, explaining its presence in adjacent non-carcinomatous epithelium. In the same sample, the significant anchor with the second highest effect size (0.348) reports on differential expression of keratin genes *KRT16* and *KRT17* (Supplementary Figure 2A); the carcinoma predominantly expresses *KRT17* while uninvolved epithelium expresses mainly *KRT16*. Both *KRT16* and *KRT17* are inducible keratins and can be expressed in squamous cell carcinoma (Huang et al. 2019; Moll, Divo, and Langbein 2008).

Applied to human fetal intestine, the anchor with the highest effect size corresponds to known alternative splicing of *RPS24* (Olivieri et al. 2021) where a 3-nt microexon is included in epithelial cells and excluded in stromal cells (Supplement Figure 2B). In the main electric organ tissue of electric eel, sc-SPLASH identifies another aspect of *RPS24* alternative splicing: electrocytes include exon 6 while stromal cells in the insulating septa exclude exon 6 (Figure 1F). This is evolutionarily consistent: *RPS24* exons are homologous between eel and human; electrocytes are derived from the skeletal muscle lineage (Gotter, Kaetzel, and Dedman 1998) and skeletal muscle is one of the few human cell types that exclude exon 6 (Olivieri et al. 2021). Together, these findings show that sc-SPLASH can be extended to spatial transcriptomics discovery, and could readily be adapted to future new barcoded sequencing technologies.

Genomic references are incomplete; in nonmodel organisms, references might be especially poor, even those constructed with state-of-the-art long reads. Alignment can therefore limit the scope of analysis. To demonstrate this, we applied sc-SPLASH to a 10x Chromium dataset from the freshwater sponge *Spongilla lacustris* (Musser et al. 2021). The 27-nt anchor GCCATCAGAACCCCAGGAACCATCTAA (which we call the granny anchor’) with the highest entropy (6.2; associated with 667 distinct target sequences; Figure 2A) was absent from both the reference genome assembly (odSpoLacu1.1) and the NCBI nucleotide database (no match by BLAST) and had relatively high effect size, consistent with cell-type specific expression. Post-facto analysis revealed that the granny anchor was predominantly expressed in granulocytes (63%, 47 out of 74 granulocytes); there is also lower expression in a minority of cells of other cell types, notably amoebocytes (Figure 2B); both granulocytes and amoebocytes are thought to be involved in immune defense (Musser et al. 2021). Granulocytes accounted for 64% of all annotated cells expressing the granny anchor. Two-color RNA-FISH for the granny anchor and the granulocyte marker ACP5 show that 88% of ACP5-positive cells co-express both; occasional cells are granny-positive and ACP5-negative (Figure 2C).

**Figure 2.**
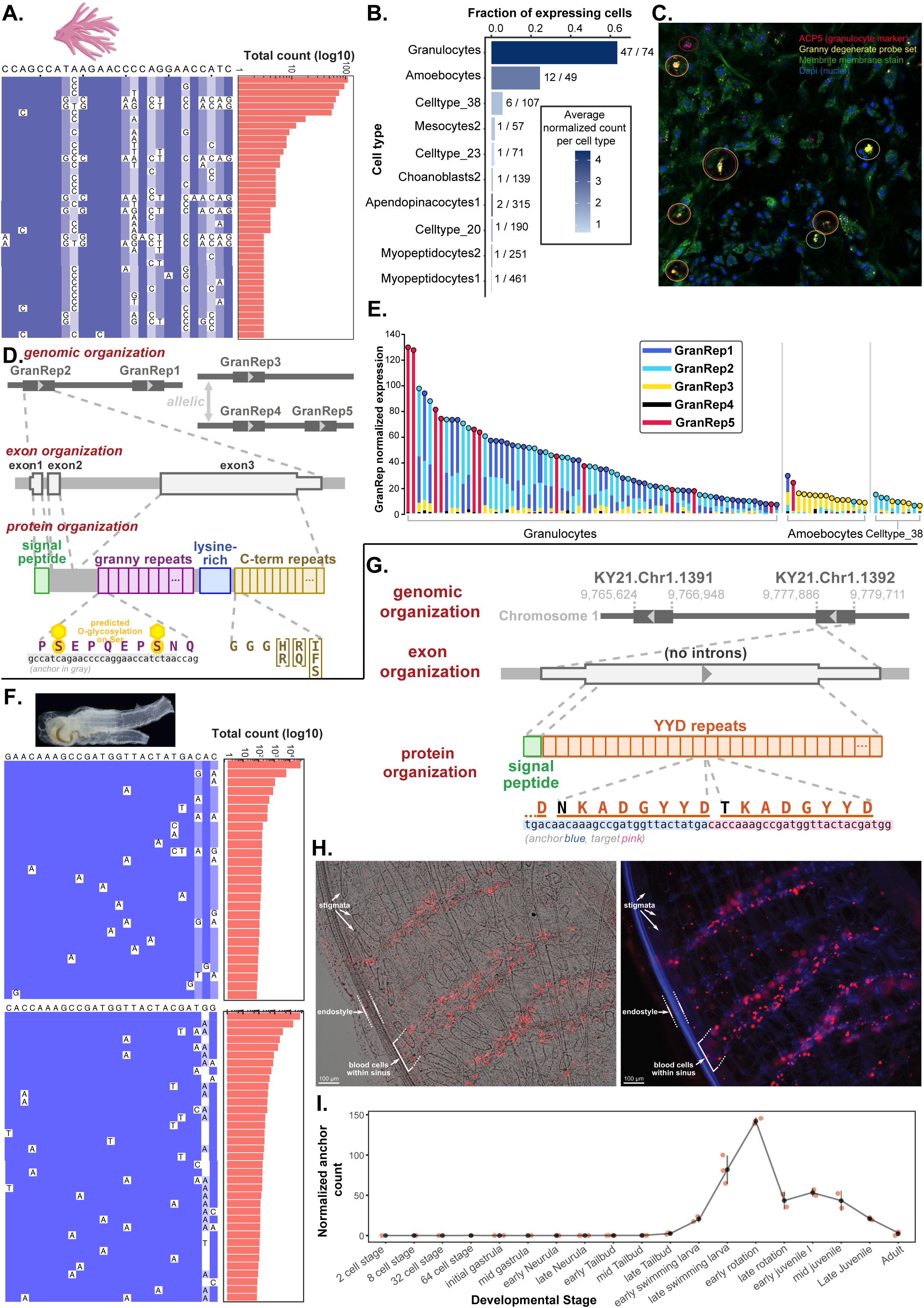
*Spongilla* and *Ciona* repeat genes with target diversity are differentially expressed. **A.** Multiple sequence alignment (MSA) of Spongilla granny anchor targets shows high target sequence diversity. The plot includes those with at least three reads across the sponge 10x dataset. **B.** Number and fraction of cells per celltype expressing the granny anchor (X/Y = expressing/total cells per celltype), suggesting predominant expression of the anchor in granulocytes and amoebocytes. Bars are colored by average normalized granny anchor count, calculated per cell as anchor count/UMI count×10^5^. **C.** HCR RNA-FISH confirms granny anchor expression in granulocytes, where cells co-expressing *ACP5* a granulocyte marker (red) and a probe set designed against different granny versions (yellow). **D.** Genomic structure of the GranRep gene family: GranRep1 and GranRep2, as well as GranRep4 and GranRep5, are on the same contig. All genes share the same 3-exon structure with granny repeats in exon 3, encoding a signal peptide, granny repeat region (30-bp repeats), lysine-rich region, and C-terminal repeats (18-bp repeats). Repeat numbers and region sizes vary by gene. **E.** Single-cell differential expression of GranRep genes suggesting granulocytes primarily express GranRep1/GranRep2, while amoebocytes primarily express GranRep3. Normalized expression per cell is calculated as aligned reads/UMI count×10^5^. GranReps are ordered by abundance in each stack, with marker colors showing the most abundant gene. Cells with ≥10 GranRep reads and normalized expression ≥5 are shown. **F.** MSA of target sequences for *Ciona* YYD anchor suggesting substantial target diversity for this anchor. We show the targets most similar to the two found in the HT genomic reference with the highest counts in the dataset. **G.** Two genes in the HT genome are composed almost entirely of YYD repeats, except for a signal peptide. **H.** HCR RNA-FISH at the juvenile stage shows YYD anchor expression restricted to circulating hemocytes. Red channel (Cy5) in both images is FISH for YYD repeat; left is a merge with brightfield, right is a merge with DNA stain (blue channel, DAPI). **I.** YYD anchor expression across Ciona development peaks during metamorphosis (“early rotation”). Normalized count per sample is calculated as anchor count/total reads×10^5^.

Subsequently, we analyzed long-read datasets to determine the sequence contexts of the granny anchor, and identified a 30-nt granny repeat in a family of five genes, that we named GranRep1 to GranRep5 (Figure 2D). Genes GranRep1 and GranRep2 are linked (on the same genomic contig); GranRep4 and GranRep5 are linked and are allelic to GranRep3. The encoded proteins are predicted to be secreted, with a similar structure: N-terminal signal peptide, 10-amino acid granny repeats (linked to the anchor), a lysine-rich region, and 6-amino acid C-terminal repeats; the granny repeats are predicted to be O-glycosylated (Figure 2D). From just four sponge individuals we found two to six alleles of each gene (Supplement). There is differential expression within the GranReps: granulocytes divide into two subsets, one subset expressing nearly exclusively GranRep5, and the other expressing mainly a combination of GranRep1 and GrapRep2; amoebocytes predominantly express GranRep3 (Figure 2E). GranRep expression increases during development from gemmules (day 0) to day 12; in a preliminary study, there was no evident difference in expression with lipopolysaccharide or cyclic dinucleotide challenge (Supplementary Figure X).

After discovering the GranRep genes, we used sc-SPLASH to search for related phenomena. We queried several marine invertebrate organisms (methods) for anchors with high sequence diversity and with evidence of being contained in a repeat (allowing for some mismatch) found in assembled sequences extended from the anchor. The top hit was the anchor TGACAACAAAGCCGATGGTTACTATGA from 10x data of the ascidian tunicate *Ciona robusta* (also known as *Ciona intestinalis* Type A) (Cao et al. 2019). While also aquatic and filter-feeders like sponges, tunicates are phylogenetically distant from sponges, being part of phylum Chordata where they are the sister group most closely related to vertebrates. The anchor had a moderate entropy of 4.80-5.97, with dozens of distinct targets with appreciable read counts (Figure 2F). NCBI BLASTN of the anchor finds it in two *C. robusta* gene models (LOC108950787 and LOC108950295); both labeled as “pesticidal crystal protein Cry11Bb-like”, the similarity being to peptide repeats at the C-terminus of *Bacillus thuringiensis* Cry11Bb protein rather than to the folded domains common to crystal toxins (Orduz et al. 1998). A more recent HT genome assembly (Satou et al. 2019, 2022) has two adjacent genes (KY21.Chr1.1391 and KY21.Chr1.1392) with eleven and nine copies of the anchor respectively, and six distinct targets in total (Figure 2G). Thus the genome assembly does not account for the observed diversity in the targets of this anchor. Examination of long-reads used in the genome assembly did not find evidence of missed anchor-containing genes. Some of the observed diversity is likely due to allelic variation: *C. robusta* has a haplotypic diversity rate of ∼1.1% (Satou et al. 2012), and gene models from genome assemblies from different individuals differ in repeat number and sequence. Like the GranReps, both encoded proteins are predicted to be secreted; the mature proteins are nearly entirely composed of imperfect repeats, with a quite different amino acid composition than GranReps (Figure 2G). We name them “YYD” repeats, after part of the amino acid sequence.

The 10x data has samples from early embryo to larva stage; anchor expression is detected at the late tail-bud stage and more prevalent in larva; although found in multiple annotated cell types (predominantly from mesenchyme, designated only by number), unannotated cells (1,221) form the largest single category. We performed RNA-FISH for the anchor at the juvenile stage (which follows larva and metamorphosis stages) when most mature organs have formed. In the juvenile, anchor expression is limited to a portion of circulating hemocytes within the hemolymph vessels (Figure 2H). Multiple types of *Ciona* hemocytes are implicated in immune defense (Longo et al. 2021) -- a striking parallel to the expression of *Spongilla* GranReps in granulocytes and amoebocytes. It is unclear if anchor expression at the larval stage is also due to hemocyte-like cells, or there is a shift in cell type over development. Anchor counts from a *C. robusta* developmental time-course dataset (bulk RNA-Seq) (Hu et al. 2017), show a peak anchor expression during early metamorphosis (“early rotation”) (Figure 2I). An interesting possibility is that immune cells may be needed to deal with the massive cell death that occurs during metamorphosis (Parrinello, Cammarata, and Parrinello 2018). Considering together the *Spongilla* and *Ciona* genes, a general hypothesis might be that repeats can be a way to rapidly create new proteins, important for quickly evolving situations like immune defense.

We note that sc-SPLASH also re-discovered *Ciona trans-*splicing, which pervasively modifies transcripts in this organism as it does in others, e.g., *C. elegans* (Lasda and Blumenthal 2011) as the most diverse anchor in the data was GAGTACATGGG<u>ATTCTATTTGAATAAG</u>, the final 16 bp (underlined) are the *Ciona* spliced leader (SL) sequence (Satou et al. 2006; Matsumoto et al. 2010), and the 5’ portion is an adaptor. This anchor had 142,486 different targets, 1,324 of which had at least 0.01% of total reads for this anchor.

## Conclusion

sc-SPLASH is an ultra-efficient statistical approach for uncovering RNA regulation, without relying on alignment. While developed for barcoded scRNA-seq, it can also be applied to barcoded spatial transcriptomics to find spatially regulated RNA variation. We demonstrated its versatility across a range of organisms, detecting sequence variations missed by reference alignment, including in spatial transcriptomics such as human tumors. It identified highly abundant secreted repeat proteins putatively expressed in immune cells and possibly markers of sub-specialization in two disparate animals–a sponge and a tunicate–illustrating new biology missed by reference-first approaches. This biological discovery represents just a glimpse of the potential for insights in barcoded sequencing data achievable with a statistics-first framework, particularly for non-model organisms.

## Acknowledgments

J.S. is supported by the National Institute of General Medical Sciences grant R35 GM139517 and the Chan Zuckerberg Data Insights. M.K. and S.D. are supported by the National Science Center, Poland (project no. DEC-2022/45/B/ST6/03032). Sponge sequencing data was generated at The Yale Center for Genome Analysis which is in part funded by the National Institutes of Health grant 1S10OD030363-01A1.

## Methods

**File downloads:**

- Human T2T assembly was downloaded from:

https://www.ncbi.nlm.nih.gov/datasets/genome/GCF_009914755.1/

- The sequences for germline V, D, and J human immunoglobulin genes sequences were downloaded from the IMGT website:

https://www.imgt.org/download/V-QUEST/IMGT_V-QUEST_reference_directory/Homo_sapien s/IG/

### Data Availability

The FASTQ files for the Tabula Sapiens data were downloaded from https://tabula-sapiens-portal.ds.czbiohub.org. The FASTQ files for Visium datasets were downloaded from the SRA database: human small intestine (SRP284357), human cutaneous squamous cell (SRP262989), electric eel muscle (SRP432489) (samples P4_ST_vis_rep1 and P4_ST_vis_rep2) were downloaded from the SRA with accession IDs and, respectively. Celltype annotations for Tabula Sapiens dataset were downloaded from figshare: https://figshare.com/articles/dataset/Tabula_Sapiens_release_1_0/14267219. The FASTQ files for Ciona datasets were downloaded from the SRA database: scRNA-seq 10x (SRP198321) and developmental timecourse data (SRP339256 and DRP003810) *Spongilla* PacBio genomic data from the Musser lab is accession # XXX; PacBio IsoSeq data is # XXX; Illumina data is # XXX. *Spongilla* PacBio genomic data from Edmonton, Canada is SRA Project PRJEB58939.

### Software Availability

sc-SPLASH is integrated into SPLASH2 pipeline and can be accessed on GitHub: https://github.com/refresh-bio/splash

BKC module for preprocessing and k-mer counting in 10x is available on GitHub: https://github.com/refresh-bio/bkc

### Hardware

Run time and memory consumption were evaluated on a machine equipped with AMD 3995WX 64-Cores CPU, 512 GB RAM, and 4 HDDs configured in RAID5.

### SPLASH framework

SPLASH directly takes FASTQ files and parses sequencing reads to extract *anchors*, or specific *k*-mers that are followed by a set of diverse *k*-mers, called as *targets*. Anchors with multiple associated targets can be used to characterize a wide range of sequence variations including mutations, alternative splicing, V(D)J recombination, RNA editing, etc. SPLASH’s core objective is to identify anchors with sample-dependent target sequence distribution where the notion of “sample” can encompass diverse biological contexts such as cells, individuals, tissues, etc. In the context of 10x scRNA-seq and Visium spatial transcriptomic, each single cell and each spatial spot, distinguished by a unique barcode sequence, is considered as a “sample”, respectively. For each anchor, SPLASH builds a contingency table of counts with rows and columns representing targets and samples, respectively. SPLASH then performs a computationally efficient statistical test that yields a closed-form p-value under the null hypothesis that target frequencies for each sample come from the same distribution. SPLASH splits the samples in two disjoint groups and the effect size for each anchor ranging from 0 (the same target distribution between two sample groups) to 1 (disjoint targets between two sample groups). Finally, anchors with a multiple testing corrected p-value <0.05 are identified as having a significant, sample-dependent target distribution.

### STAGE 1 of sc-SPLASH - BKC

Stage 1 of sc-SPLASH is devised from scratch to add support for barcoded reads. To this end, we designed BKC (barcoded-reads *k*-mer counter), a specialized *k*-mer counting tool tailored for barcoded format. BKC is executed separately for each sample, but several BKC instances can be executed in parallel (if there are many samples).

Each BKC run’s input is a set of FASTQ file pairs. In each FASTQ pair, the reads are treated together: the _1.fastq file contains barcode and UMI, while _2.fastq contains cDNA. In the first step, BKC works only on _1.fastq files. It loads barcode+UMI reads and builds in memory a dictionary containing the tuples <barcode, UMI, file_id, read_id>. Both barcode and UMI are packed and are represented as 64-bit integers. Each input file is processed in parallel, and a separate dictionary is prepared for it.

Then, the file-oriented dictionaries are merged into a single dictionary. This dictionary is pruned. There are several options here. When the user provides a white list of barcodes, we treat them as *trusted*. Otherwise, we determine a threshold between *trusted* and *non-trusted* barcodes in the same way as in UMI-tools, i.e., we look for the knee in the curve showing the cumulative number of reads per barcode. Optionally, the non-trusted barcodes can be corrected if a single symbol substitution allows them to be changed uniquely into a trusted one. Then, we remove the dictionary entries with non-trusted barcodes. Finally, we do UMI deduplication. We only allow a unique dictionary entry for barcode+UMI pair. The selection of the preserved read is random but deterministic.

In the second step of BKC, we deal with _2.fastq files. Initially, we load the reads into memory, for which entries are in the pruned dictionary built in the previous step. To save space, we store them using 3-to-1 packing (3 bases are represented in a single byte, which allows us to distinguish A, C, G, T, and N symbols and the end-of-read marker). The loading is performed in parallel, and the number of threads can be up to the number of input files. Then, depending on the mode, we count *k*-mers or *k*-mer pairs. We do this in parallel using the user-given number of threads. Each thread handles reads for some subset of barcodes and does the following. First, it enumerates all *k*-mer pairs (or *k*-mers) in all reads for the currently processed barcode. Then, it sorts them and gathers statistics. Optionally, various filters can be applied here, e.g., removal of polyACGT runs, removal of artifacts (like Illumina adapters or any other given by the user), and removal of rare *k*-mers (according to the user-provided criteria). Finally, the gathered statistics for each barcode are compressed and stored in the output file. The compressed files can be used directly by sc-SPLASH in further stages. They can also be dumped into a textual format for other pipelines.

BKC can also work in the filter mode, in which it does barcode filtering and UMI deduplication of input reads. Due to parallelization and implementation in the C++ programming language, it is much more efficient than UMI-tools in this mode.

### STAGE 2 of sc-SPLASH

It was also necessary to redesign the second stage of SPLASH2 to add support for barcodes. Now we have to load all records from each bin. This could potentially increase the memory footprint, but in practice, this component is not memory-dominating. Then we sort the records according to: anchor+target+sample_id+barcode. For each unique anchor, we construct a contingency table, in which columns are for sample_id+barcode and rows are for target. Then, we proceed in the same way as in SPLASH2.

### STAGE 3 of sc-SPLASH

The final stage of sc-SPLASH is the same as in SPLASH2.

### Performance evaluation against Cell Ranger and STARsolo

We compared the running time and memory requirements of sc-SPLASH v2.11.3 against STARsolo v2.7.10b and Cell Ranger v8.0.1 on TSP1 and TSP2 datasets. For each run, we used 16 threads. In the case of STARSolo and Cell Ranger it is required to run each sample separately. For TSP1, the summary running time/memory usage peak across all samples was 7622s/70GB, 1476s/35GB, and 316s/8GB for Cell Ranger, STARSolo, and sc-SPLASH, respectively. For TSP2 it was 11195s/64GB, 2673s/35GB, and 510s/18GB accordingly. In the case of sc-SPLASH it is also possible to run all samples at once. In this case, the running time/memory usage was 137s/22GB and 399s/40GB for TSP1, and TSP2, respectively.

We used the following command lines:

~~~
cellranger count --id=$sample_id --fastqs=$data_path --sample=$sample_name
--localcores 16 --create-bam true
--transcriptome=/home/refdata-gex-GRCh38-2020-A

STAR --genomeDir /data/T2T_human_genome/STAR_index_files --readFilesIn
$input_files --soloType CB_UMI_Simple --runThreadN 16 –soloCBwhitelist
3M-february-2018.txt --soloUMIlen 12 --readFilesCommand gunzip -c

splash --technology 10x --anchor_len 27 --target_len 27
--n_threads_stage_1 1 --n_threads_stage_1_internal 16 --n_threads_stage_2
16 --without_compactors input.txt
~~~

In the case of running all samples at once with sc-SPLASH we used the following command line:

~~~
splash --technology 10x --anchor_len 27 --target_len 27
--n_threads_stage_1 4 --n_threads_stage_1_internal 4 --n_threads_stage_2
16 --without_compactors input.txt
~~~

### Performance evaluation of BKC against UMI-tools

To evaluate the performance of BKC 1.1.0 for input reads filtering against UMI-tools 1.1.6 we used TSP1_S13 dataset (2 runs, 98.86 GB in total). BKC needed 126s and 7GB RAM to determine trusted barcodes, barcode filtering, and UMI deduplication. The command line was:

~~~
./bkc --mode filter --input_name fl --export_filtered_input_mode both --n_threads 8 --verbose 1 --apply_cbc_correction
~~~

UMI needed 5360s (about 40x longer time) and 0.6 GB RAM for determining trusted barcodes and barcode filtering. We were not able to do UMI deduplication as UMI-tools is used in alignment-based

pipelines, and UMI deduplication is made after mapping. The command lines were:

~~~
umi_tools whitelist --stdin=r1.fastq.gz
   --bc-pattern=CCCCCCCCCCCCCCCCNNNNNNNNNNNN --log=whitelist.log
   --stdout=whitelist.tsv

umi_tools extract --bc-pattern=CCCCCCCCCCCCCCCCNNNNNNNNNNNN
   --whitelist=whitelist.tsv \
   --stdin=TSP1_Muscle_Abdomen_10X3primev31_1_1_S13_L003_R1_001.fastq.gz \
   --stdout=L3_1_filtered.fastq.gz \
   --read2-in=TSP1_Muscle_Abdomen_10X3primev31_1_1_S13_L003_R2_001.fastq.gz \
   --read2-out=L3_2_filtered.fastq.gz

umi_tools extract --bc-pattern=CCCCCCCCCCCCCCCCNNNNNNNNNNNN
   --whitelist=whitelist.tsv \
   --stdin=TSP1_Muscle_Abdomen_10X3primev31_1_1_S13_L004_R1_001.fastq.gz \
   --stdout=L4_1_filtered.fastq.gz \
   --read2-in=TSP1_Muscle_Abdomen_10X3primev31_1_1_S13_L004_R2_001.fastq.gz \
   --read2-out=L4_2_filtered.fastq.gz
~~~

We note that BKC filtering has differences from UMI-tools. BKC keeps only a single read for each UMI+cell barcode; UMI-tools use of mapping information may allow more sophisticated UMI deduplication.

### Running SPLASH on scRNA-seq datasets

SPLASH was run on the data from each individual and tissue in Tabula Sapiens separately. Fastq files from different lanes of the same 10x library were run together to allow UMI deduplication across different lanes of the same library. Both anchor and target lengths were set to 27 with no gap between each anchor and target. Cells with >2000 UMIs were retained for statistical analysis. Other SPLASH parameters were set as default, namely, anchors that have at least 50 reads in total and are found in at least 2 cells are kept for analysis. For each cell, only anchors with at least 5 reads are considered. Prior to statistical analysis, any anchor with A/G/C/T stretches of at least 7bps or a 15-bp match with the UniVec database, containing vector, adapter, or primer sequences commonly used for cDNA sequencing, was filtered out. We used a cutoff of 0.05 for p-values after the Benjamini-Yekutieli correction and an effect size threshold of 0.2 to call anchors.

### Calculating entropy for target diversity quantification

For each anchor, we compute the target entropy as a statistical measure of target sequence diversity. Assume that the anchor *A* has *T* targets across *N* samples. Let *t_i_* (1 ≤ *i* ≤ *T*) be the marginal count across all samples for target *i*, i.e., if *t* denotes the target count for target *i* in sample *j*, then 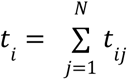. We compute entropy *H_A_* for anchor *A* as: 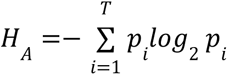, where 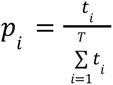 fraction of reads belonging to target *i*. For extreme cases where all samples express the same target sequence and each sample expresses a separate target (*T* = *N*) with the same count (*t_i_* = *t*, 1 ≤ *i* ≤ *T*), the target entropy would be 0 and *log*_2_ *T*, respectively.

### Alignment of SPLASH calls to reference genome

While sc-SPLASH’s inference is entirely reference-free, to enhance the interpretability of the results, we also implemented a *post facto* alignment step. For each pair of a significant anchor and one of its targets (top 4 abundant targets in our study), we construct an *extendor* by concatenating their sequences. These extendors are built from the most abundant targets, ensuring they are free from sequencing errors and artifacts. We then align extendors to the reference genome using both STAR (Dobin et al. 2013) and Bowtie2 (Langmead and Salzberg 2012) and report their alignment information. We also assign a gene name to each extendor sequence based on its alignment position and if the reference transcriptome is also available, we provide information about whether the splice sites are annotated exon boundaries and splice junctions are annotated. For each anchor, we also compute and report the hamming distance, levenstein distance, and the longest common subsequence (lcs) distance. These metrics offer further insights into the underlying mechanism leading to target sequence diversity, even in the absence of alignment information. For example, having the same hamming distance and Levenstein distance for the targets of an anchor suggests that targets are most likely due to SNPs or mutations, as the sequence variations between targets can be attributed solely to substitutions.

### Running IgBLAST

We ran IgBLAST (Ye et al. 2013) through the `AssignGenes.py` function in Change-O toolkit (Gupta et al. 2015). We selected the sequences for IgBlast that were unaligned or were aligned by STAR to an immunoglobulin gene. After running IgBlast, those sequences with an annotated immunoglobulin locus (IGH, IGK, IGL) that were also in frame and productive were selected as the V(D)J recombination sequences.

### Pfam analysis

We performed a homology-based search for the extendors for each significant anchor to obtain annotated protein domains from the Pfam database (Finn et al. 2016). For each sequence, 6 possible reading frames (3 for each strand) are used for in silico translation, and if the q-value for the Pfam best Pfam annotation across all reading frames is <0.01, it is reported for the extendor.

### Supervised test for identifying metadata-dependent calls

SPLASH performs unsupervised inference in the sense that it does not need class membership information about the input samples which is usually available as metadata files. In the context of scRNA-seq, the supervised step is especially useful for finding cell type-specific RNA sequence diversity. We formulate this as a regression problem where we aim to find anchors whose target distribution can predict the metadata group of the samples (cells). Let C_i be the metadata class for the ith sample (cell), and t_ij denotes the count for target i in sample j, we consider the following multinomial logistic regression:

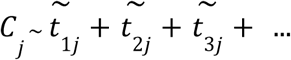

Where *t̃*_1*j*_ denotes the fraction of reads in sample j that belongs to target 1. We perform an L1-regularized regression as implemented in GLMnet (Friedman, Hastie, and Tibshirani 2010), and report those anchors with nonzero regression coefficient as metadata-dependent calls (or cell type-specific in case of scRNA-seq analysis).

### Running SPLASH on Visium datasets

We ran SPLASH on each Visium slide separately. If the Visium slide had more than one FASTQ file, all of them were run through SPLASH together. We downloaded their spatial metadata files: `tissue_hires_image.png, scalefactors_json.json,` and `tissue_positions_list.csv` from the NCBI GEO database. We considered only those spots whose barcodes were identified as within-tissue in `tissue_positions_list.csv.`

### Sponge culture

Adult sponges with gemmules in the overwinter stage were collected from the underside of a ferry dock in Lake Constance near Kressbronn, Germany in February 2022 and stored at 4°C until used for experiments. Juvenile sponges (*Spongilla lacustris*) were grown by placing gemmules in culture dishes (Falcon #353001) with M-medium (1 mM CaCl_2_·6H_2_O, 0.5 mM MgSO_4_·7H_2_O, 0.5 mM NaHCO_3_, 0.05 mM KCl, 0.25 mM Na_2_SiO_3_) (Rasmont 1961), and kept at 18°C in the dark. M-medium was replaced every other day, beginning around day 5 when juvenile sponges first adhere to the cover slip.

### Sponge developmental time course

To survey DNA and RNA variability across development, we sampled two replicates of approximately 25 juvenile sponges at each of days 5, 8, and 12 after plating gemmules in M-medium. These days correspond to developmental stages 3, 4, and 5, respectively (Funayama et al. 2005). DNA and RNA were extracted from juvenile sponges at each stage using the Qiagen DNA/RNA Allprep Micro Kit (Qiagen #80284), and the resulting extracts were quantified using a Nanodrop 1000 and Qubit. DNA integrity was determined using an Agilent TapeStation to ensure sufficient quality (DNA Integrity Number > 6). For one replicate of each stage we obtained less than 100 ng of total DNA and for these performed 4 PCR cycles using the Equinox Library Amplification Kit (Watchmaker Genomics #7K0014-096) following kit instructions. RNA sample integrity was determined on Bioanalyzer to ensure high quality (RNA integrity number > 8).

### Sponge stimulation of immune response

To determine changes in DNA and RNA sequences and abundance following stimulation of an immune response, we performed treatments with either lipopolysaccharides (LPS) from *Pseudomonas aeruginosa* (Sigma #L9143) or 2′3′-cGAMP (Invivogen #tlrl-nacga23-02) in M-medium. LPS treatments were conducted in dishes containing five sponges 7-days after gemmule plating (stage 3) by replacing M-medium with premixed 5ug/uL LPS with M-medium. This concentration was chosen based on unpublished results indicating it induces a significant immune response in freshwater sponges (Scott Nichols, pers. comm.). Juvenile sponges were then harvested for DNA and RNA either one day later (day 8, stage 4) or had their LPS treatment terminated at day 8 by replacing LPS M-medium with fresh M-medium, followed by harvesting at day 12 (stage 5). Treatment with cyclic dinucleotides was performed for four hours in 8-day old (stage 4) and 12-day old (stage 5) sponges by replacing M-medium with premixed 100uM 2’,3’-cGMP in M-medium. DNA and RNA were isolated following similar protocols as for our developmental time course samples, resulting in high-quality DNA from two replicates and RNA from one replicate for each treatment and time point.

### Sponge DNA and RNA sequencing

DNA samples were purified using Ampure XP spri beads (Beckman Coulter # A63882) and library preparation was performed using the Watchmakers Library Prep Kit with Fragmentation (Watchmaker Genomics #7K0013-096). RNA polyA library preparation was performed using the KAPA mRNA HyperPrep Kit (Roche #KK8581). Both DNA and RNA samples were sequenced at the Yale Center for Genome Analysis on a Novaseq X Plus at 2×150bp to obtain 100 million reads per sample.

### Sponge in-situ hybridization chain reaction

To validate the expression of granny in specific sponge cell types, we performed in-situ Hybridization Chain Reaction (HCR) experiments on stage 5 juvenile sponges exhibiting all adult features, including osculum, a well-developed canal system, and numerous choanocyte digestive chambers. Sponges were labeled with Membrite-488 (Biotium #30093) and fixed in 4% paraformaldehyde in ¼ strength Holtfreter’s Buffer (10x HF buffer = 590mM NaCl; 6.7mM KCl; 7.6mM CaCl_2_; 2.4mM NaHCO). The HCR protocol from Scott Nichols (Nichols 2023) was followed. Minor changes included no RNase Inhibitor treatment, probe hybridization with 1uM probe solutions, amplification with 6pmol of hairpins, and counterstaining with Hoescht (Thermo #H1398) before mounting in Fluoromount G (Southern Biotech #0100-01) for imaging. Probe sequences are listed in the supplementary data. Images of the HCR samples were taken on the LSM 880 Airyscan NLO/FCS Confocal Microscope, A Zeiss Axio Observer Z1 inverted microscope, using Zen 2.1 software. Over two experiments, 30 cells were ACP5+ granny+, 4 were ACP5+ granny–, and 4 were ACP5– granny+.

### Sponge local DNA assembly, expression quantification, glycosylation prediction

GranRep genomic contigs were assembled manually by sequential overlap extension of PacBio HiFi reads. All reads containing the granny anchor (by `grep`) were collected and aligned with Clustal Omega. The multiple sequence alignment (MSA) was manually divided into clusters; the GranRep genes are numbered roughly by the abundance of reads in each cluster. Clusters were extended by `grep` for *k*-mers (*k* ≤ 50) near the ends of the cluster, represented in at least two reads. If `grep` gave a large number of read hits (indicating match to a repetitive element), another *k*-mer was chosen. The new reads were aligned with the previous round reads. Extension of GranRep1 reached GranRep2; extension of GranRep4 reached GranRep5. In some rounds for GranRep1, the new MSA showed obviously two clusters (due to structural variation); in this case, reads were split into separate clusters and extension was applied to each. We also reviewed reads for single-nucleotide variation, and together with structural variation, were able to infer two complete haplotypes for GranRep1, one of which contains a ∼10 kb mobile element insertion with a highly repetitive internal sequence. The exon structures were established by correlation with RNA-Seq. The resulting gene models (from PacBio data from the Musser lab) were used as reference transcripts for Bowtie alignment and expression quantitation in the 10x data. In a few cases, variant transcript models were assembled from Illumina paired-end RNA-Seq using a combination of Bowtie mapping to the above reference transcripts and `grep` extension. Contigs and transcript models are deposited at NCBI, with accession numbers XXXX. O-glycosylation was predicted with NetOGlyc-4.0 (Steentoft et al. 2013).

### *Ciona* In-situ Hybridization Chain Reaction

Short in situ hybridization antisense DNA probes were designed based on the split-probe design of HCR v.3.0 using HCR 3.0 Probe Maker (Elagoz et al. 2022; Kuehn et al. 2022) with adjacent B1 amplification sequence. 20 probe pairs were designed and ordered as oligo pools (Integrated DNA Technology) and suspended in nuclease-free water at a concentration of 0.5 µM. Probe sequences are listed in the supplementary data.

For in situ hybridization, *C. robusta* juveniles were incubated in fixation buffer (1× phosphate buffered saline (PBS) and 4% paraformaldehyde) overnight at 4 °C. Fixed samples were then dehydrated in methanol and stored at −20 °C for at least 24 h and up to several months. The samples were progressively rehydrated in PBS. They were permeabilized in detergent solution (1.0% SDS, 0.5% Tween-20, 150 mM NaCl, 1 mM EDTA (pH 8), 50 mM Tris-HCl at pH 7.5) for 30 min. The samples were extensively washed in PBSX1 and then in 5× saline sodium citrate buffer containing 0.1% Tween-20 (SSCT), before being prehybridized in hybridization buffer (Molecular Instruments) for 30 min at 37 °C. The probes were then added to the hybridization buffer at a final concentration of 0.05 µM and the samples were allowed to hybridize at 37 °C overnight. Following hybridization, the samples were washed three times for 30 min in probe wash buffer (Molecular instruments) at 37 °C and then in 5× SSCT at room temperature. They were then pre-amplified in amplification buffer (Molecular Instruments) for 30 min. Meanwhile, H1 and H2 components of the HCR hairpins B1 coupled to Alexa647 fluorophores (Molecular Instruments) were incubated separately at 95 °C for 90 s, cooled down to room temperature in the dark and then pooled together before being added to the amplification buffer at a final concentration of 60 nM. The amplification was then performed overnight at room temperature. The samples were subsequently washed three times for 30 min in 5× SSCT and incubated in PBS containing 1.6:1,000 DAPI (Invitrogen) for 1 h. Samples were mounted in 50% glycerol diluted in PBSX1 for imaging.

The specificity of the antisense DNA probes and amplification hairpins was validated by running the protocol without probes.

### List of invertebrates queried for repeat sequences with high diversity

We queried the 10x scRNA-seq datasets from the following marine invertebrates for finding anchors with high diversity that are contained within a repeat sequence: *berghia stephanieae* (sea slug), *biomphalaria glabrata* (freshwater snail), *ciona robusta* (sea squirt), *loligo vulgaris* (common squid), octopus, *exaiptasia diaphana* (sea anemone aiptasia), soft coral Xenia, *Nematostella vectensis* (starlet sea anemone), and sea urchin.

## Notes

### Competing Interest Statement

The authors have declared no competing interest.

